# Coregistered transcranial optoacoustic and magnetic resonance angiography of the human brain

**DOI:** 10.1101/2022.05.24.492943

**Authors:** Ruiqing Ni, Xose Luis Dean-Ben, Valarie Treyer, Anton Gietl, Christoph Hock, Jan Klohs, Roger M. Nitsch, Daniel Razansky

## Abstract

Imaging modalities capable of visualizing the human brain have led to major advances in neurology and brain research. Multi-spectral optoacoustic tomography (MSOT) has gained importance for studying cerebral function in rodent models due to its unique capability to map changes in multiple hemodynamic parameters and to directly visualize neural activity within the brain. The technique further provides molecular imaging capabilities that can facilitate early disease diagnosis and treatment monitoring. However, transcranial imaging of the human brain is hampered by acoustic attenuation and other distortions introduced by the skull. Here, we demonstrate noninvasive transcranial MSOT angiography through the temporal bone of an adult healthy volunteer. Time-of-flight (TOF) magnetic resonance angiography (MRA) and T_1_-weighted structural MRI were further acquired to facilitate anatomical registration and interpretation. The superior middle cerebral vein in the temporal cortex was identified in the MSOT images, matching its location observed in the TOF-MRA images. These initial results pave the way toward the application of MSOT in clinical brain imaging.

## TEXT

Research into brain functionality and neurodegenerative disorders critically depends on the availability of bioimaging tools capable of noninvasive observations with high resolution and specific contrast [1, 2]. Functional magnetic resonance imaging (fMRI) has enabled a better understanding of brain function and connectivity abnormalities in brain diseases, becoming a workhorse in neuroimaging [3]. Nuclear imaging technologies, such as positron emission tomography (PET) and single-photon emission computed tomography (SPECT) assisted with radiolabelled tracers, have further enabled the molecular diagnosis and monitoring of neurodegenerative diseases such as Alzheimer’s disease (AD) and Parkinson’s disease [4, 5]. However, these well-established technologies are commonly associated with high installation and maintenance costs, use of ionizing radiation and/or limited accessibility. These drawbacks have fostered the development of new imaging approaches that can complement or enhance imaging performance and provide better affordability and operability. For example, functional near-infrared spectroscopy (fNIRS) capitalizes on optical contrast to measure changes in both oxygenated (HbO_2_) and deoxygenated hemoglobin (Hb), which is not possible with fMRI [6, 7]. However, strong light diffusion in living tissues severely limits the achievable resolution. Functional ultrasound (US) provides highly sensitive imaging of cerebral blood flow and has been used in humans, e.g., through the temporal bone, despite the strong acoustic distortions induced by the skull [8]. In addition, magnetoencephalography (MEG) systems [9-11] and transcranial ultrafast US localization microscopy [12] have been developed to better map brain function and microcirculation.

Multi-spectral optoacoustic tomography (MSOT) provides unique capabilities for visualizing murine brain activity, including high spatio-temporal resolution and functional specificity [13]. In particular, its five-dimensional (real-time spectroscopic three-dimensional) imaging capability enabled new insights into large-scale neuronal activity and the accompanying hemodynamic changes [14-16]. Moreover, MSOT imaging of optical probes targeting amyloid-beta deposits and tau fibrils has enabled visualizing specific accumulation of aggregates in animal models of AD, multiparametric characterization of glioblastoma tumors, and neuroinflammation in ischemic stroke [17-24], which may serve to define new diagnostic biomarkers. MSOT imaging, however, faces similar transcranial imaging challenges as fUS. The relatively thin murine skull bone has been shown to induce acoustic aberrations in broadband optoacoustic (OA) signals, particularly in the high frequency components, leading to minor distortions and loss of contrast and resolution of the images [25]. Much more significant acoustic distortions are induced by the thick human skull, which further supports complex non-linear propagation patterns involving longitudinal to shear mode conversion and guided acoustic waves [26, 27]. Efforts have been devoted to the development of new algorithms trying to correct for skull-induced aberrations [28-31], for which data from other modalities such as X-ray computed tomography (CT) and MRI can be used [32, 33]. More recently, MSOT has been shown to provide powerful capabilities to map brain activity matching fMRI readings in patients who underwent hemicraniectomy [34]. However, no transcranial MSOT imaging of human subjects has thus far been achieved.

In this letter, we demonstrate the basic feasibility of visualizing cerebral vasculature through the temporal bone in humans with transcranial MSOT imaging. The results were validated and registered with head-to-head time-of-flight (TOF) magnetic resonance angiography (MRA) and T1-weighted structural MRI. The imaging system consisted of a spherical matrix array (Imasonic SaS, Voray, France) accommodating 256 piezocomposite elements and covering an angle of 90° [35]. All the elements have an approximate size of 3×3 mm^2^, central frequency of 4 MHz and detection bandwidth of ∼100%, providing an almost isotropic resolution of 200 μm in a region close to the center of the spherical array geometry. The excitation light was provided with an optical parametric oscillator (OPO) laser tunable between 680 and 1250 nm and delivering ∼40 mJ energy per pulse at a pulse repetition frequency (PRF) up to 100 Hz (Innolas, GmbH, Krailling, Germany). The optoacoustically generated signals were digitized at 40 megasamples per second with a custom-designed data acquisition system (Falkenstein Mikrosysteme GmbH, Taufkirchen, Germany) triggered with the Q-switched output of the laser and transmitted to a personal computer via 1 Gbit/s ethernet.

Noninvasive MSOT imaging of the brain of a healthy female volunteer was performed through the temporal bone with the spherical array operated in a hand-held mode. For this, a custom-designed holder 3D-printed in polylactic acid (PLA) was attached to the array and filled with agar (1.3% agar powder w/v) to guarantee acoustic coupling [36]. The laser beam was guided through a fiber bundle with 5 output arms (**Fig. 1a**). The laser wavelength was hopped between 700, 800 and 1064 nm at a pulse repetition frequency of 25 Hz. The laser fluence at the skin surface was ∼12 mJ/cm^2^, i.e., well below safety thresholds for human exposure [37]. The probe was gently scanned along the left temporal bone. During the image acquisition, the eyelids were closed and further protected with black tape to avoid any potential unwanted exposure. Image reconstruction was performed with a graphics processing unit (GPU) implementation of a back-projection algorithm [38]. This reconstruction method enabled real-time rendering to preview the images during acquisition, thus facilitating the localization of the structures of interest. Spectral un-mixing was performed with a standard least-square fitting approach to the spectra of oxygenated and deoxygenated hemoglobin after normalizing the signals with the wavelength-dependent laser fluence [36]. All processing steps were performed in MATLAB (MathWorks, Natick, MA, USA). Individual 3D images were then compounded [39] to provide a larger field-of-view spanning 30×30 mm in the lateral dimension.

**Fig 1.**
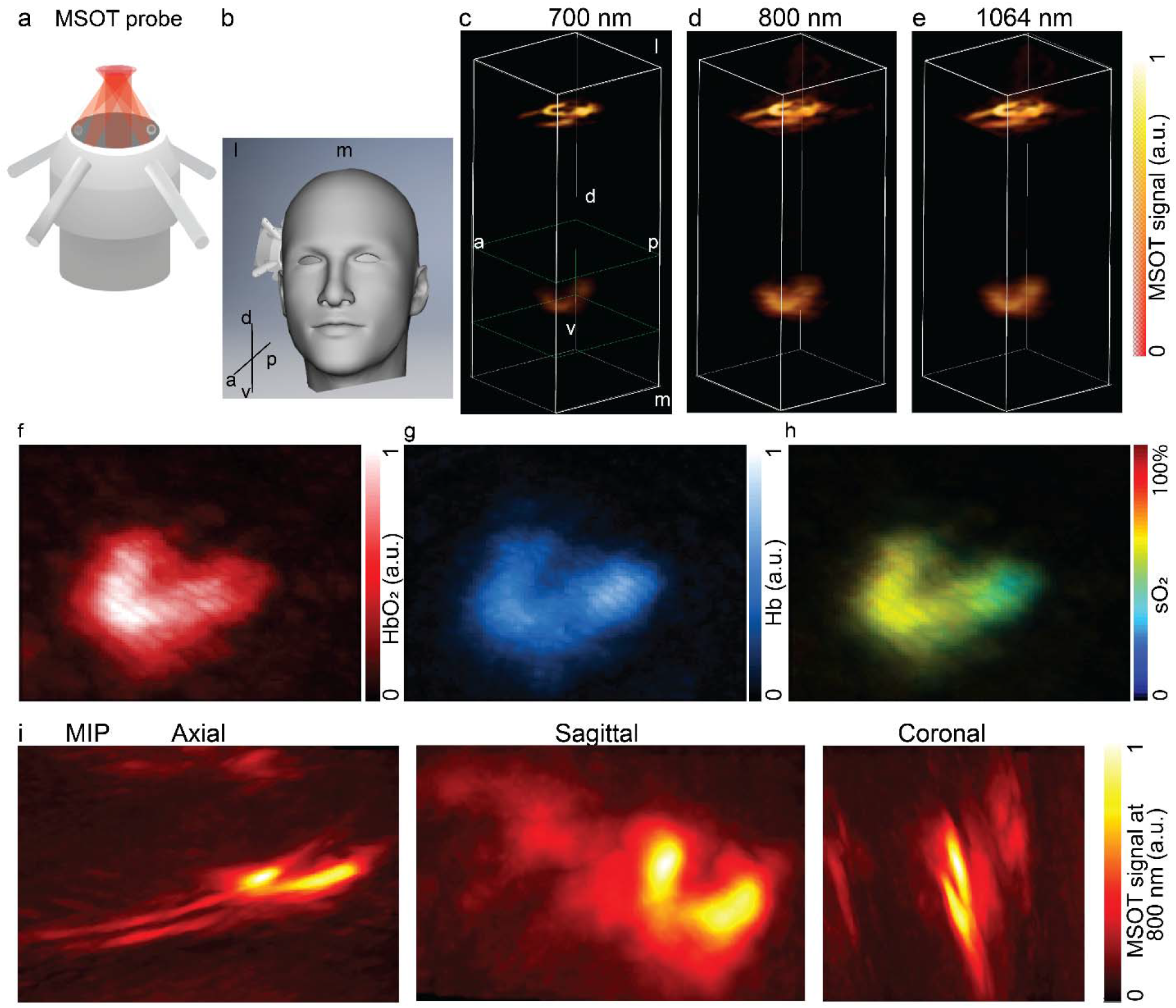
Transcranial MSOT imaging of the human brain through the temporal bone using a hand-held probe. (**a**) Illustration of the multifiber hand-held MOST imaging probe (diameter 6 cm). The laser beam was guided through a fiber bundle with 5 output arms to facilitate a more uniform illumination pattern on the skin surface. Beam diameter at the tissue surface was approx. 15 mm. (**b**) MSOT imaging procedure through the temporal bone using the hand-held probe. (**c-e**) MSOT images acquired at 700, 800 and 1064 nm. Field-of-view 10×10×25 mm. (**f-h**) Spectrally unmixed distributions of hemoglobin (HbO_2_) and deoxyhemoglobin (Hb) and the resulting oxygen saturation (sO_2_) of the deep vessel inside the brain (indicated by green box in c). (**i**) MSOT image recorded at 800 nm - maximum intensity projections (MIPs) are shown in the axial, sagittal, and coronal orientations. m, medial; l, lateral; a, anterior; p, posterior; d, dorsal; v, ventral.

*In-human* transcranial brain images acquired with MSOT through the temporal bone are shown in **Fig. 1b**. The effective imaging depth was sufficient to clearly visualize the temporal cortex and superior middle cerebral vein (SMCV), also known as the Sylvian vein, located at an approximate depth of 13 mm from the skin surface (lower parts of **Figs. 2a-c**). This vessel was identified as a vein based on an oxygen saturation (sO_2_) value in the of 60-70% range (**Figs. 1f-h**), whilst sO_2_∼100% is usually expected in arteries. The deep vessel SMCV was more prominent at 800 nm and 1064 nm, arguably due to the higher light attenuation at shorter wavelengths. The superior temporal vein (STV) could also be localized (upper parts in **Figs. 1c-e**), with more prominent contrast manifested at 700 nm. The wavelength-dependent signal patterns in deep tissue differed from the more superficial signals generated in the skin/muscle, indicating that the vascular patterns in the brain were not associated with skull reflections.

**Fig 2.**
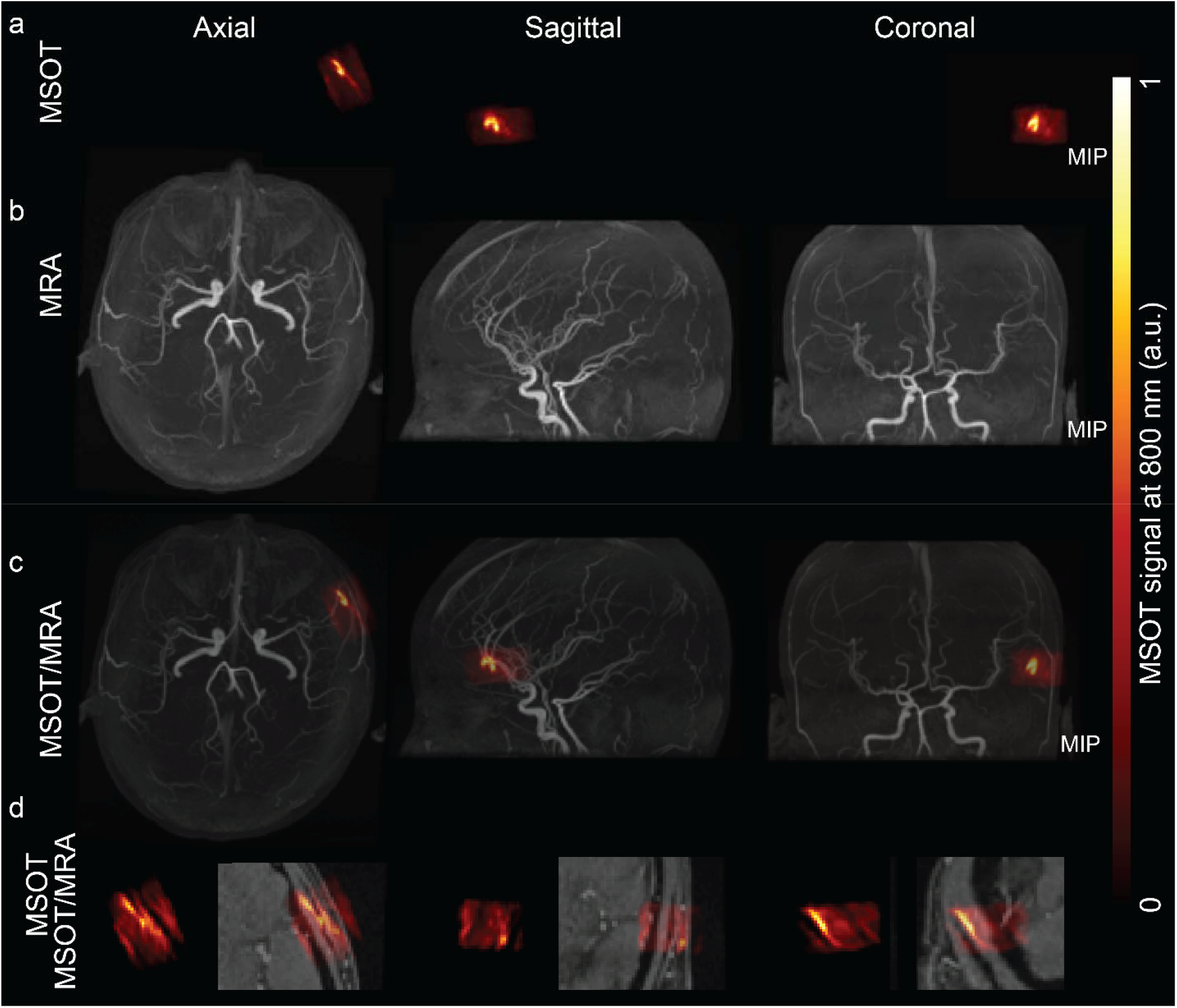
Registration of MSOT and magnetic resonance angiography (MRA) of human brain. (**a**) MSOT image recorded at 800 nm - maximum intensity projections (MIPs) are shown in the axial, coronal and sagittal orientations of the temporal cortex region; (**b**) Time-of-flight (TOF)-MRA MIPs are shown in axial, coronal and sagittal view; and (**c**) Overlay of **a** and **b** in axial, coronal, and sagittal view. (**d**) zoomed-in of MSOT image recorded at 800 nm and overlay of MSOT image recorded at 800 nm overlay with TOF-MRA in axial, coronal and sagittal view;

To map the exact thickness of the temporal bone and location of the STV and deeper cerebral vessels, TOF-MRA and T_1_-weighted MRI scans were performed in the same subject at University Hospital Zurich using a GE Signa Premier 3T scanner (**Fig. 2, 3**). Standard clinical 3D TOF MRA sequences (slice thickness = 1.2 mm, repetition time (TR) = 21 ms, time to echo (TE) = 3.4 ms, flip angle = 20 degrees, autocalibrating reconstruction for Cartesian imaging) with HyperSense acquisition acceleration and a 48-channel head coil provided optimal vascular contrast [40]. A native T_1_-weighted MR magnetization-prepared 180 degrees radio-frequency pulse and rapid gradient-echo (MP RAGE) were used for structural reference (slice thickness = 1 mm, TR = 2191.36 ms, TE = 2.996 ms, inversion recovery = 900, flip angle = 8 degrees, spacing = 0.5, 512×512×312 image matrix, 0.4688×0.4688×0.5 mm voxel size) [41]. PMOD software (PMOD Technologies GmbH, Zurich, Switzerland) was subsequently used for segmentation and registration of the MRI and MSOT data. An overlay of the MSOT and MRA data (**Fig. 1)** provided validation of the location of the SMCV in the temporal cortex in the MSOT images. The SMCV and STV vessels measured approximately 1 mm in diameter, as seen from the TOF MRI (**Fig. 2, 3, SVideo1**). While similar STV dimensions were rendered with both modalities, the SMCV appears smeared in the MSOT images (**Fig. 1**), which is arguably attributed to severe acoustic aberrations introduced by the skull. The skull thickness was estimated to be approximately 2-3 mm in the temporal bone area in the T_1_-weighted images.

**Figure 3.**
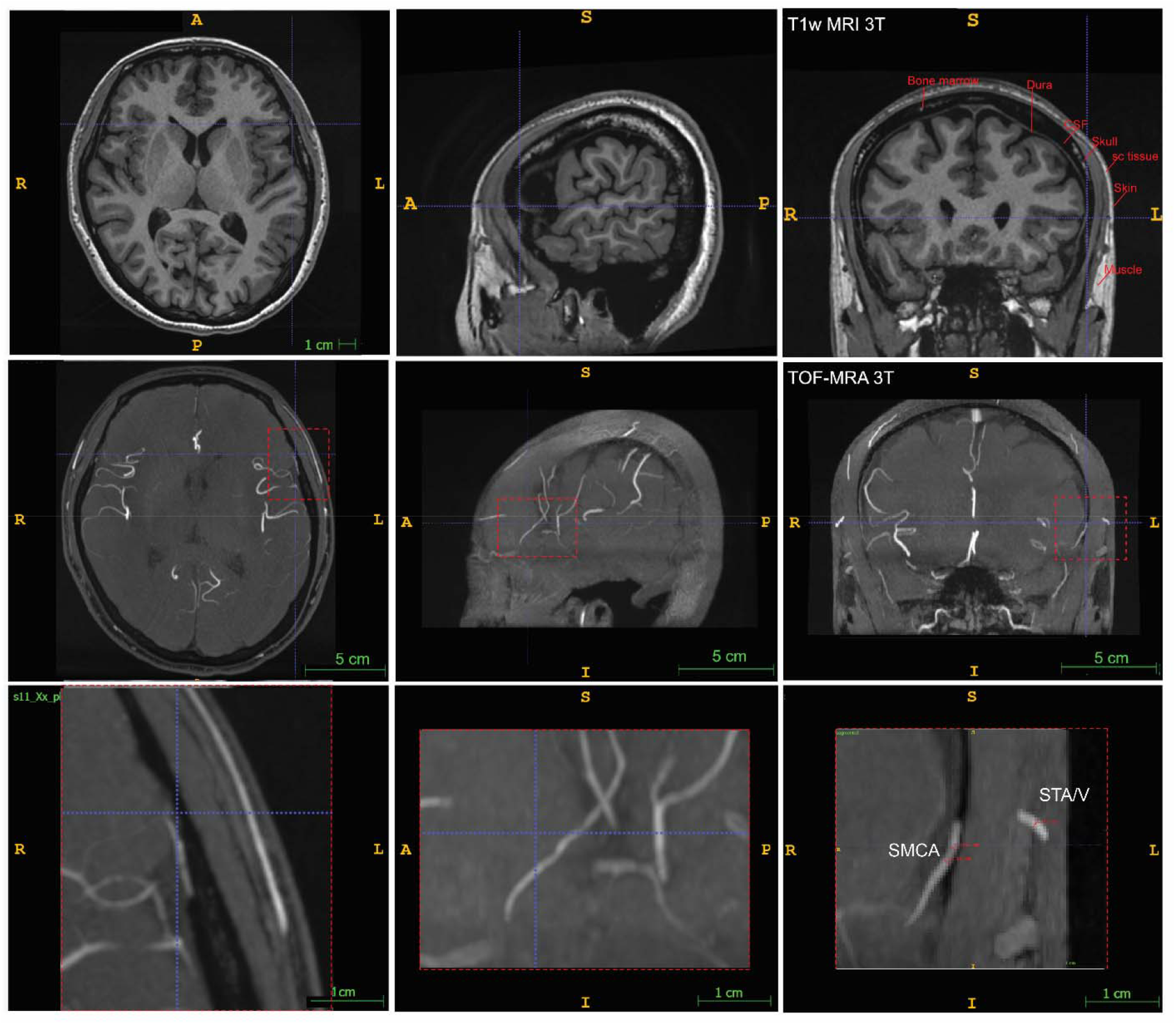
T_1_-weighted MRI and TOF-MRA images indicating the location and diameter of the vessels. r, right; l, left; a, anterior; p, posterior; d, dorsal; v, ventral. superior middle cerebral vein (SMCV), superior temporal vein (STV).

This work demonstrates the basic feasibility of performing noninvasive MSOT brain angiography through the temporal bone in humans. Our results indicate that human skull induces strong acoustic aberrations resulting in significant distortion of deep vascular structures. We further investigated to what extent the images were afflicted with other potential sources of image distortions. In particular, pressure waves generated within subcutaneous blood vessels undergoing acoustic reflections in the skull may result in spurious structures mirrored into the brain. However, in our experiments, no strong contrast was generated at the skin surface that could potentially result in strong reflections. Furthermore, the vascular structures in the MSOT images acquired at different wavelengths exhibited a distinctive absorption spectrum. Finally, TOF-MRA images also served to validate the actual location of the vessels identified in the MSOT images. We are thus confident that the observed structures actually correspond to the pial vasculature.

MSOT imaging is based on tissue excitation with nonionizing NIR radiation that is safe for human use. The laser energy density employed in this work was below safety standards causing no thermal or other adverse effects. Several experimental aspects can be further optimized. For example, the efficiency of excitation light delivery can be improved based on a photon recycler that has been shown to recover ∼50% of the optical energy loss [42]. US attenuation is generally known to increase with frequency [43], e.g., resulting in ∼80% signal loss at 1 MHz through the full skull thickness [44], which makes it easier for the low frequency signal components to traverse the skull. Nevertheless, skull transmission windows are known to exist at certain frequencies [45]. The diminished thickness of the temporal bone facilitates US transmission, which has previously been exploited for pulse-echo US imaging of the middle cerebral artery (MCA), the anterior cerebral artery (ACA) and the posterior cerebral artery (PCA) [46]. Likewise, acoustic sensors may be positioned within the naval cavity or ocular regions, which have been shown to facilitate US access to the brain [32].

Apart from causing strong acoustic dispersion and attenuation, the high acoustic impedance of the skull bone (∼7.7 × 10^6^ Rayls [47]) relative to the soft brain tissue results in loss of spatial resolution when assuming a uniform speed of sound for the MSOT reconstruction procedure. Accurate modelling of transcranial US propagation is not straightforward, as it implies taking into account mode conversion at surfaces and guided wave propagation [27]. The development of reconstruction algorithms accounting for these effects could potentially be facilitated with accurate prior anatomical information available in the MRI images. Indeed, the feasibility of hybridization between preclinical MSOT and MRI has recently been demonstrated [48, 49]. A method for correcting the distortion in pulse-echo transcranial US images acquired through the temporal bone has recently been proposed based on estimating aberration delays from the backscattered US waves in individual microbubbles [12]. Combining MSOT and pulse-echo US [50-52] may then further enable correcting for acoustic distortions in the MSOT images. Additionally, absorbing microparticles could also be localized and tracked individually in real time with MSOT [53]. These and similar particle localization approaches may also facilitate the development of dedicated MSOT reconstruction algorithms tailored for transcranial imaging, e.g., based on the optoacoustic memory effect [54].

The MSOT capacity for transcranial imaging in humans can potentially define new application niches in clinical brain imaging. MSOT offers a unique capability to visualize multiple hemodynamic parameters associated with brain activity [14]. The recent validation of MSOT readings in humans with clinical fMRI [55] anticipates the great potential of MSOT to study brain function in health and disease. The temporal cortex is involved in cognitive functions, e.g., memory, perception and language, and is affected in many brain disorders, including AD [66], frontotemporal dementia [67], major depressive disorder [68], and temporal lobe epilepsy [69]. MSOT has recently demonstrated high molecular sensitivity for targeting various brain diseases in preclinical models and can potentially be used for defining new biomarkers for *in vivo* clinical brain research and diagnosis.

## Supporting information

SVideo 1

## Funding

RN received funding from Helmut Horten Stiftung and University of Zurich [MEDEF-20-021]. DR acknowledges funding from the US National Institutes of Health grants UF1-NS107680 and RF1-NS126102.

## Disclosures

The authors declare no conflicts of interest.

## Notes

### Competing Interest Statement

The authors have declared no competing interest.

